# Cancer relevance of human genes

**DOI:** 10.1101/2021.02.04.429823

**Authors:** Tao Qing, Hussein Mohsen, Vincent L. Cannataro, Michal Marczyk, Mariya Rozenblit, Julia Foldi, Michael F. Murray, Jeffrey P. Townsend, Yuval Kluger, Mark Gerstein, Lajos Pusztai

**Affiliations:** Breast Medical Oncology, School of Medicine, Yale University, New Haven, CT, USA; Computational Biology and Bioinformatics Program, Yale University, New Haven, CT, USA; Department of Biology, Emmanuel College, Boston, MA, USA; Department of Data Science and Engineering, Silesian University of Technology, Gliwice, Poland; Department of Genetics, Yale Center for Genomic Health, New Haven, CT, USA; Department of Biostatistics, Yale School of Public Health, New Haven, CT, USA; Department of Pathology, School of Medicine, Yale University, New Haven, CT, USA; Applied Mathematics Program, Yale University, New Haven, CT, USA; Department of Molecular Biophysics & Biochemistry, Yale University, New Haven, CT, USA; Department of Computer Science, Yale University, New Haven, CT, USA; Department of Statistics & Data Science, Yale University, New Haven, CT, USA

**Author notes:** Correspondence to Lajos Pusztai, MD, DPhil, Breast Medical Oncology, Yale Cancer Center, Yale School of Medicine, 300 George St, Suite 120, Rm 133, New Haven, CT, 06520, USA. These authors contributed equally (Tao Qing, Hussein Mohsen).

**Keywords:** Cancer genes, protein-protein interaction, selection pressure, effect size

## Abstract

**Background:** It is unclear how many of genes contribute to the biology of cancer. We hypothesize that genes that interact with core cancer gene (CCG) in a protein-protein interaction network (PPI) may have functional importance.

**Methods:** We categorized genes into 1- (n=6791), 2- (n=7724), 3- (n=1587), and >3-steps (n=362) removed from the nearest CCG in the STRING PPI and demonstrate that the cancer-biology related functional contribution of the genes in these different neighborhood categories decreases as their distance from the CCGs increases.

**Results:** Genes closer to cancer genes manifest greater connectedness in the network, show greater importance in maintaining cell viability in a broad range of cancer cells in vitro, are also under greater negative germline selection pressure in the healthy populations, and have higher somatic mutation frequency and cancer effect.

**Conclusions:** Approximately 70% of human genes are 1 or 2 steps removed from cancer genes in protein network and show functional importance in cancer-biology. These results suggest that the universe of cancer-relevant genes extends to thousands of genes that can contribute functional effects when dysregulated.

## Background

An important goal of cancer research is to identify genes that are relevant to cancer biology and decipher their functional contributions to malignant transformation. Since the discovery of the first viral oncogenes in the 1970s, alterations in hundreds of human genes have been implicated in the development of cancer through genetic association studies and in vitro and in vivo functional experiments (1–4). The terminology has also shifted, many of these genes are now referred to as “cancer driver genes” implying therapeutic potential. However, it has also become clear that even the classical transforming oncogenes are not able to transform a normal cell into a malignant cell on their own: malignant transformation requires cumulative alterations in many cellular processes often referred to as the hallmarks of cancer (5, 6). How many genes are involved in this transformation process is poorly understood. Large–scale whole-exome and whole-genome sequencing studies revealed several hundred to several thousand somatic non-synonymous single-nucleotide variants, indels, and small and large structural variants in cancers (1, 4, 7, 8). The vast majority of these somatic alterations are not recurrent in a given cancer type, appear random, and are therefore considered to be “passenger mutations” that contribute little to the biology of the disease and result from genomic instability (9). However, increasing evidence suggests that rare, non-recurrent somatic or germline alterations have functional importance and contribute to the unique clinical course of each cancer (10–12). Furthermore, a recent pan-cancer whole genome analysis found no cancer driver alterations in about 5–9% of cancers, supporting the hypothesis that the spectrum of cancer-relevant genes is broader than our current models suggest (4).

All human proteins form functional networks within cells that are connected directly through shared network members or indirectly through other proteins or functional intermediaries. For simplicity, in this manuscript we refer to genes as synonyms for their protein products. We hypothesize that proteins physically associated with, or known to directly interact with, an experimentally or clinically validated core cancer gene (CCG) can also have an impact on cancer biology and denote these genes as "one step removed" from a CCG. By extension, we also assume that genes that directly interact with the “one-step removed genes”, might also influence cancer biology, although to a lesser extent. Based on this model, one could categorize human genes into one-, two-, three-, and > three-steps removed from the nearest CCG in the network. We predict that the cancer-biology related functional contribution of the genes in these different neighborhood categories will decrease as their distance from the CCGs increases (**Figure 1**). We compared across these four mutually exclusive gene categories the average connectedness in a protein-protein interaction network, in vitro cancer cell viability scores after gene silencing, somatic mutation frequencies and cancer effect sizes in human cancers, and the negative germline selection pressure on the member genes.

**Figure 1.**
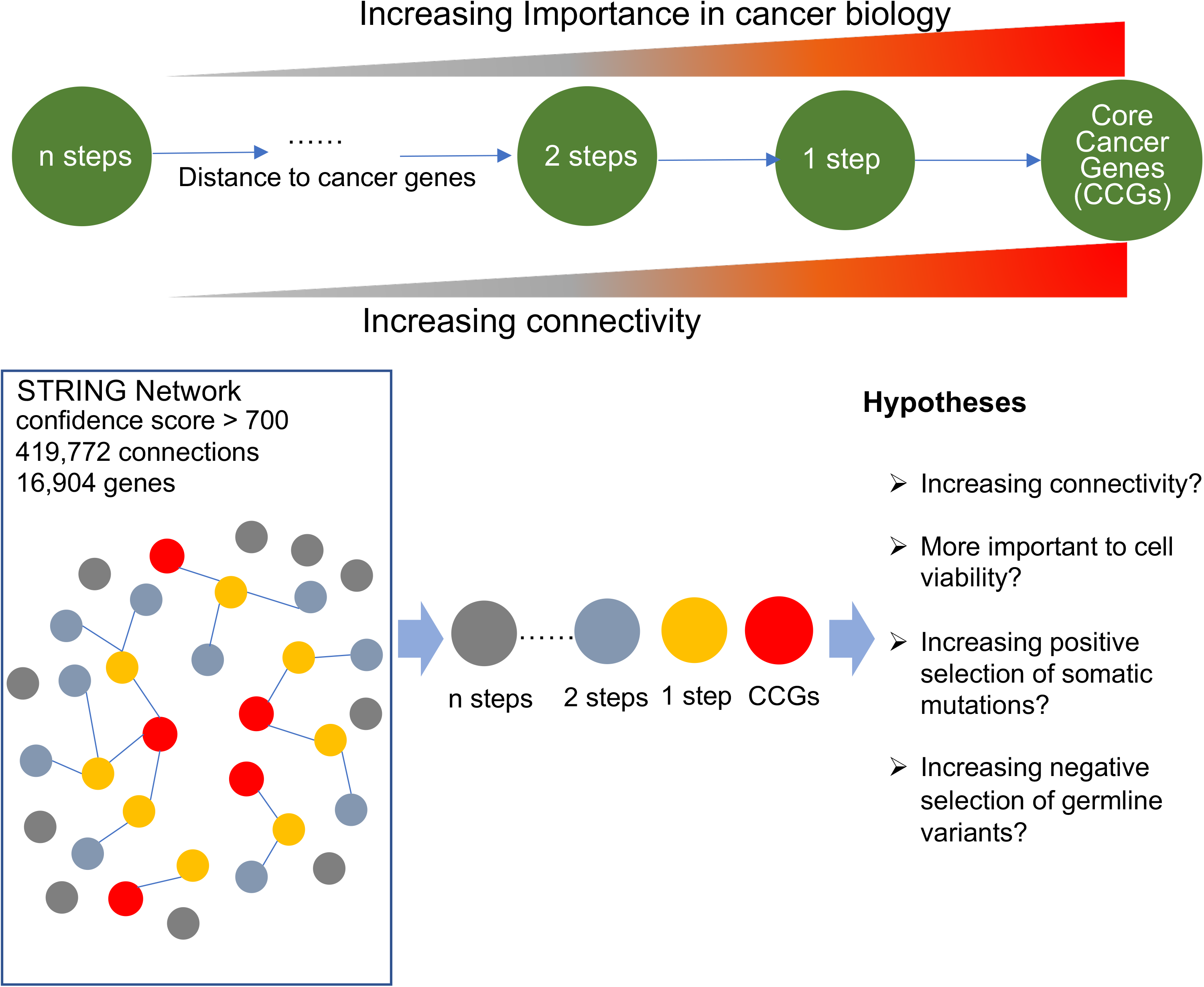
Study schema. Overview of our hypothesis that genes closer to core cancer genes in STRING network are more functional important in cancer development.

## Results

### Identify CCGs neighborhood genes in protein-protein interaction network

We designated 468 genes that comprise the Memorial Sloan Kettering-Integrated Mutation Profiling of Actionable Cancer Targets (MSK-IMPACT) as the CCGs (13, 14). Each of these genes (when their function is altered) are experimentally and clinically validated for their importance in cancer development and many also serve as therapeutic targets. We mapped these CCGs into the STRING (Search Tool for the Retrieval of Interacting Genes/Proteins) protein-protein interaction (PPI) database that includes 419,772 connections between 16,904 human proteins with a confidence score >700(15), and calculated the shortest distance (i.e. the minimum number of steps [genes]) between the CCGs and all other genes/proteins. We defined 4 gene neighborhood categories; i) genes that directly interact with CCGs (1-step removed), ii) genes that interact with CCGs through another gene (2-steps), iii) genes that interact with CCGs through 2 other intermediary genes (3-steps), and iv) genes that interact through more than 2 genes (> 3-steps) (**Figure 1, Methods**).

Four hundred forty of 468 CCGs were included in the STRING database and 6791, 7724, 1587 and 362 genes were categorized as 1 step, 2 steps, 3 steps, and >3 steps removed from the closest CCG (**Figure 2a, 2b,** Supplementary Table 1). These results indicate that most human genes are only 1 or 2 steps removed from canonical cancer genes. To find out what proportion of these cancer gene neighbors have previously been implicated in cancer biology we cross-referenced them with 2,202 genes identified as cancer drivers, oncogenes, or tumor suppressors in the CancerMine database (16). Only a small proportion—18.2%, 6.1%, 3.8% and 2.2% of the 1 step, 2 steps, 3 steps, and >3 steps neighbor genes, respectively—were linked to cancer in the literature (**Figure 2c**, Supplementary Table 1). We also calculated the network connectivity of each gene as the number of direct connections to all other genes (Supplementary Table 1). The connectivity of 16,904 genes ranged from 1 to 1,435 **(Figure 2d)**. We demonstrated that CCGs have higher connectivity than neighbor genes and gene connectivity score was gradually decreasing with increasing distance from CCGs (**Figure 2e**).

**Figure 2.**
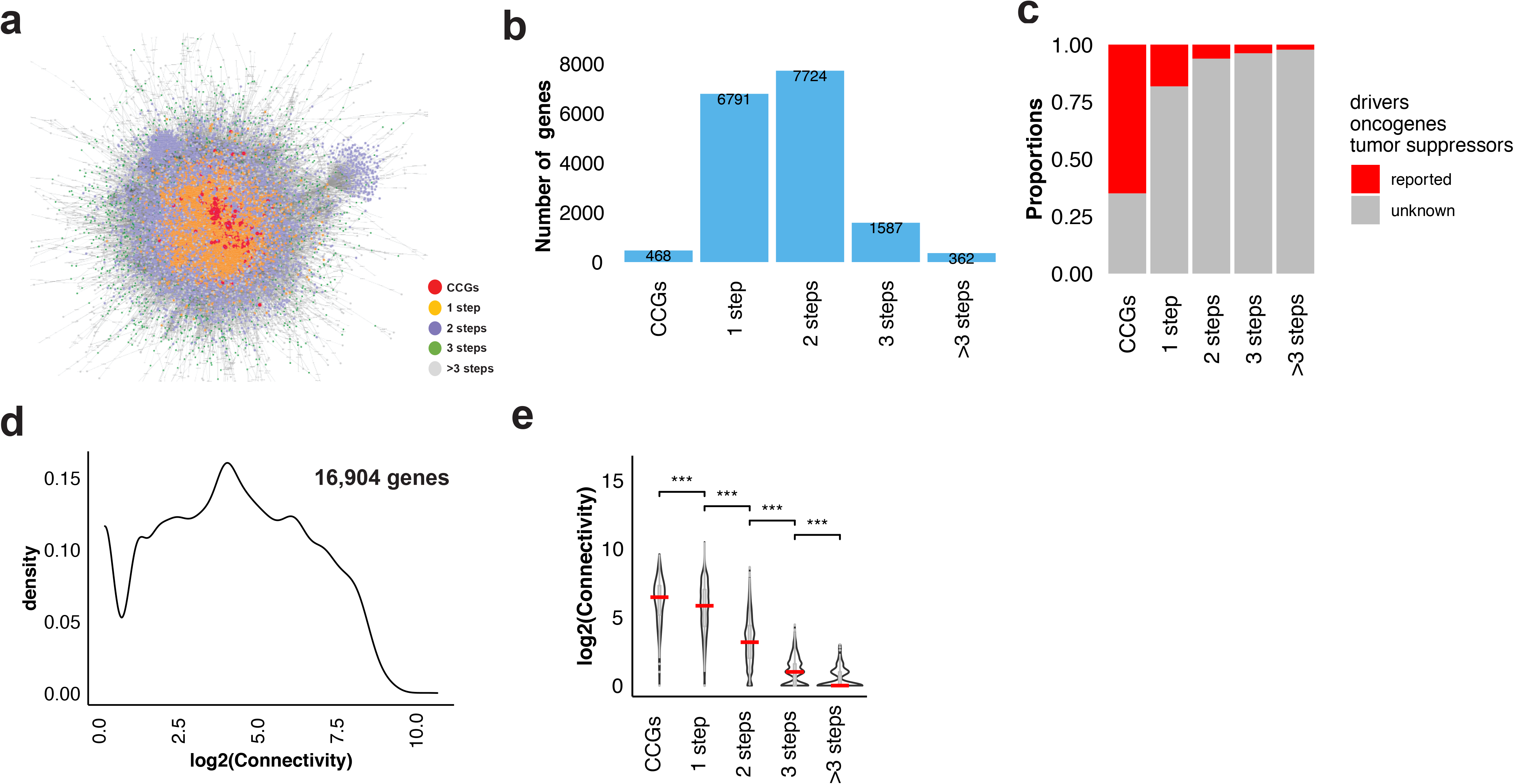
Connectedness of cancer genes. **a** STRING protein interaction network. Each dot represents a gene, colors indicate distance from core cancer genes. The grey lines show between-gene connections. **b** Number of human genes in 4 cancer gene neighborhood categories. **c** Proportion of genes implicated in cancer biology in the literature (reported or not in connection with cancer) by neighborhood categories. **d** Distribution of log_2_-transformed connectivity score of 16,904 human genes in STRING. **e** The distribution of log_2_-transformed connectivity score for the cancer genes and 4 neighborhood categories. One-sided Mann–Whitney *U* test (values of closer neighborhood genes are greater than that of all the genes in the remoter steps) *P* values are symbolized by ***, **, * corresponding to *P* < 0.0001, 0.001, and 0.01, respectively. Red bars correspond to the median of the distributions. CCGs: core cancer genes.

### The role of CCGs neighborhood genes in cancer cell viability

To estimate the functional importance of genes in the different neighborhood categories, we obtained gene dependency scores from The Cancer Dependency Map (DepMap) project. DepMap performed genome-wide pooled loss of function screening for all known human genes using RNA interference (RNAi) and CRISPR-Cas9-mediated (CRISPR) gene editing to estimate tumor cell viability after gene silencing (17). Gene dependency scores are available for 17,309 genes in 712 cell lines from shRNA, and for 17,634 genes in 563 cell lines from CRISPR experiments. A dependency score of 0 corresponds to no effect on cell viability, a negative score corresponds to impaired cell viability after knocking down the gene; the more negative the dependency score the more important the gene is for cell viability. We calculated the average dependency scores across all cell lines for the 4 categories of genes (Supplementary Table 1). CCGs had the lowest dependency scores and genes 1 step removed had statistically similar average negative dependency score, genes 2 steps and 3 steps removed had significantly negative scores closer to 0 indicating less functional importance in cell viability, and genes > 3 steps away from CCGs had average dependence scores of 0. The statistical trend for decreasing average negative dependency score with increasing distance from CCGs was statistically significant in both the CRISPR (**Figure 3a,** Kendall’s τ z-statistic = 15.13, *P* < 1 × 10^−5^) and RNAi data sets (**Figure 3b**, Kendall’s τ z = 19.33, *P* < 1 × 10^−5^). These results indicate that a very large number of human genes influence cell viability, and this influence is proportional to their distance from canonical cancer genes in the PPI network.

**Figure 3.**
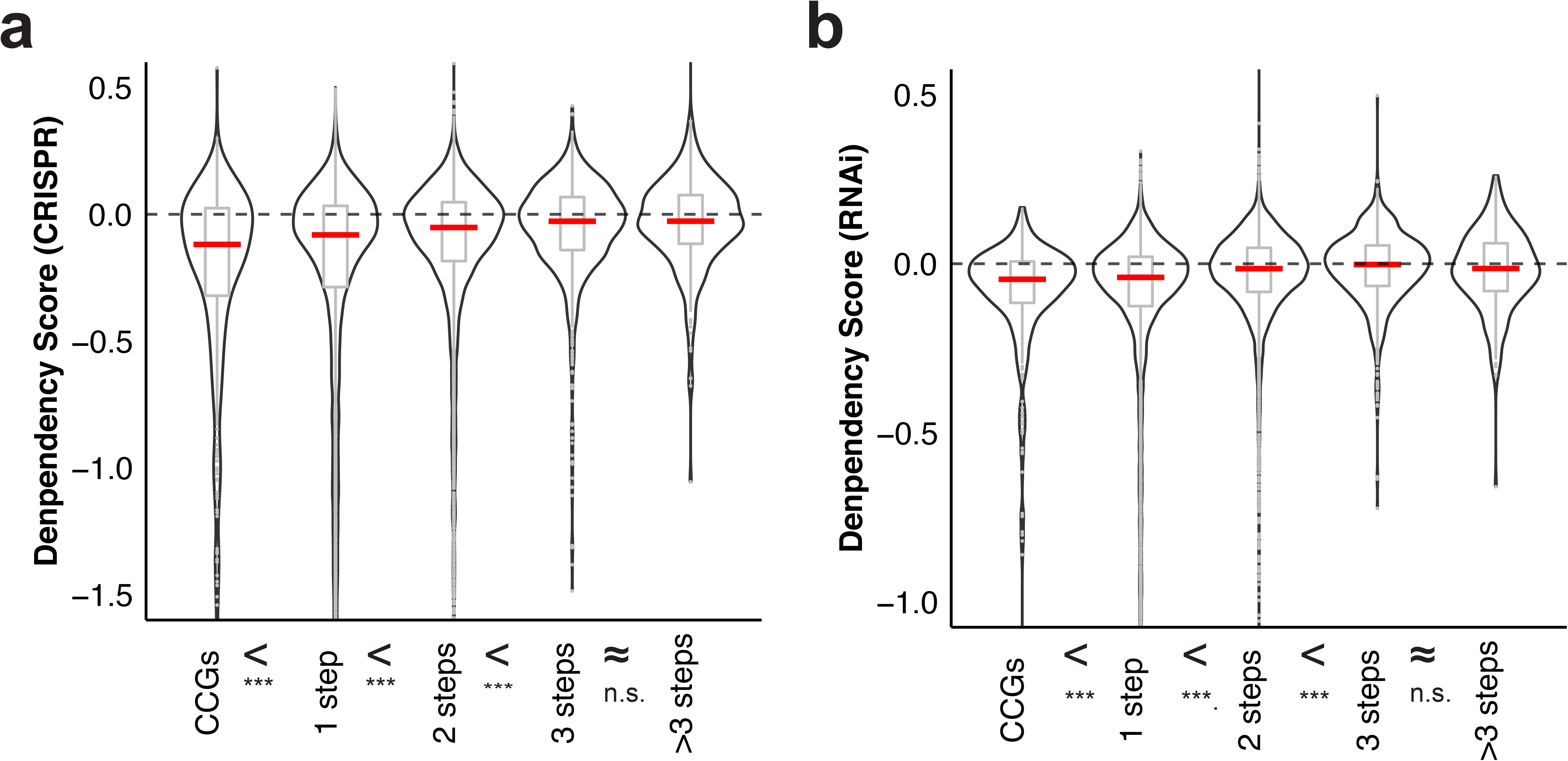
Cell viability dependence scores for cancer genes and genes in different cancer gene neighborhood categories. **a** Distribution of DepMap CRISPR-based dependency scores. **b** Distribution of DepMap RNAi-based dependency scores. *Y*-axes are dependency scores—the lower the value, the more important the gene is for cell viability. One-sided Mann–Whitney *U* test (values of closer neighborhood genes are greater than that of all the genes in the remoter steps) *P* values are symbolized by ***, **, and *, corresponding to *P* < 0.0001, *P* < 0.001, and *P* < 0.01, respectively, and n.s. abbreviating not significant. Red bars correspond to the median of the distributions. CCGs: core cancer genes.

### Positive selection of somatic mutations in CCGs neighborhood genes

The DepMap data identifies genes whose loss of function impairs cell viability. However, gain of function, inappropriate gene expression, or altered protein substrate affinity that can arise through somatic mutations can also contribute to malignant transformation. Genes whose dysfunction supports cancer growth are therefore more likely to carry somatic mutations in cancer tissues (11, 18). We compared the average prevalence of somatic mutations of CCGs and genes in the 4 neighborhood categories in 32 cancer types sequenced as part of The Cancer Genome Atlas (TCGA). Mutations of CCGs exhibited the highest average prevalence and prevalence decreased significantly with increasing distance from CCGs in 21 of 32 TCGA cancer types with substantial tumor sampling (Kendall’s τ z < −2.96, FDR < 0.018, Supplementary Tables 1 & 2; **Figure 4a**). In the remaining 11 cancer types, the prevalence of somatic mutation does not show any difference among genes in the 4 neighborhood categories (Supplementary Figure 1). Interestingly, across all cancers, the trends of somatic mutation prevalence across CCGs and neighborhood categories were negatively correlated with cancer incidence rate, tumor mutation burden, and number of somatic mutation affected genes (Supplementary Figure 2). Cancers with high incidence rate had mutations in a broader range of genes, suggesting that a larger number of genes may contribute to transformation in common cancers than in rare tumors.

**Figure 4.**
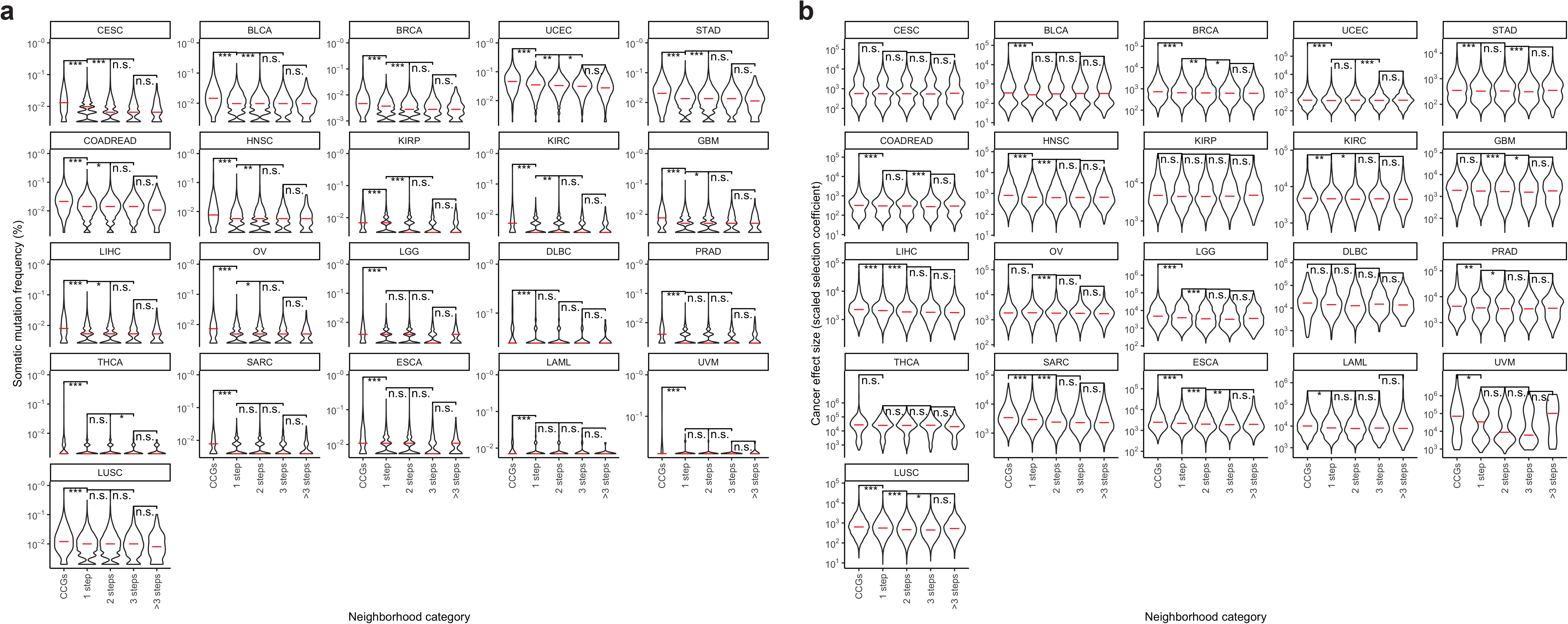
Somatic mutation frequencies of genes and cancer effect sizes of variants in genes across CCGs and 4 neighborhood categories in 21 well-sampled TCGA cancer types. **a** Somatic mutation frequencies of many TCGA types show decreasing somatic mutation frequency for genes with increasing distance from CCGs (FDR < 0.05). **b** Average cancer gene effect size (scaled selection coefficients) of variants in all genes of 4 neighborhood categories decrease with increasing distance from CCGs. Red bars correspond to the medians of the distributions. One-sided Mann–Whitney *U* test (values of closer neighborhood genes are greater than that of all the genes in the remoter steps) *P* values are symbolized by ***, **, and *, corresponding to *P* < 0.0001, *P* < 0.001, and *P* < 0.01, respectively, and n.s. abbreviating not significant. CCGs: core cancer genes.

Differences in the prevalence of mutations among these categories could arise from either increased rates of cellular mutation, or from conferring selective benefit upon cancer cell lineages. The cancer effect size is a scaled selection coefficient of the mutation, conveying the degree to which the mutation enhances the survival or reproduction of the mutant lineage(11). We further demonstrated that average effect size of single nucleotide somatic variants in all the genes of a given neighborhood category tend to decrease by the distance from CCGs in the TCGA cancer types (**Figure 4b**), suggesting that genes close to CCGs are under strong positive selections of somatic mutation which might increase cell fitness and tumor progression.

### Selection pressure of germline protein-truncating variants in CCGs neighborhood genes

Genes that play important roles in cell differentiation, cell division, cellular metabolism and that regulate physiologic cell death are also major contributors to a broad range of diseases— not just malignant transformation—when they function aberrantly. Because of their importance in maintaining normal cellular homeostasis, there is evolutionary pressure to conserve their normal DNA sequence in the germline. Deleterious germline variants in these genes decrease fitness and tend to be rare in human populations (19). The greater the functional importance, the stronger this negative selection pressure (20). Indeed, population-based whole-exome sequencing studies indicate strong negative selection pressure on deleterious germline variants in many cancer-related genes (21). We therefore hypothesized that if step-1 genes are functionally more important to cancer cell proliferation and survival than step-2 genes, which are more important than step-3 genes, we would expect to see decreasing negative germline selection pressure across these groups. Large-scale exome sequence of population genome allows to estimate the germline selection pressure for human genes in healthy individuals(20, 22). We obtained coefficients of negative selection of heterozygous rare protein-truncating variants (*S*_*h*_) for 15,998 human genes from Casa et al (20) and loss-of-function intolerance (pLI) scores for 18,225 genes from Monkel et al (22) (Supplementary Table 1). The higher the *S*_*h*_ and pLI score, the greater the selection pressure against protein-truncating germline variants of a given gene. We observed a highly statistically significant decrease in average *S_h_* (Kendall’s τ z = −27.7, *P* < 1 × 10^−5^, **Figure 5a**) and pLI (Kendall’s τ z = −29.4, *P* < 1 × 10^−5^, **Figure 5b**) scores with increasing distance from CCGs. This decrease indicates that genes closer to CCGs in the PPI network are under higher germline selection pressure, supporting greater functional importance in maintaining normal cellular functions.

**Figure 5.**
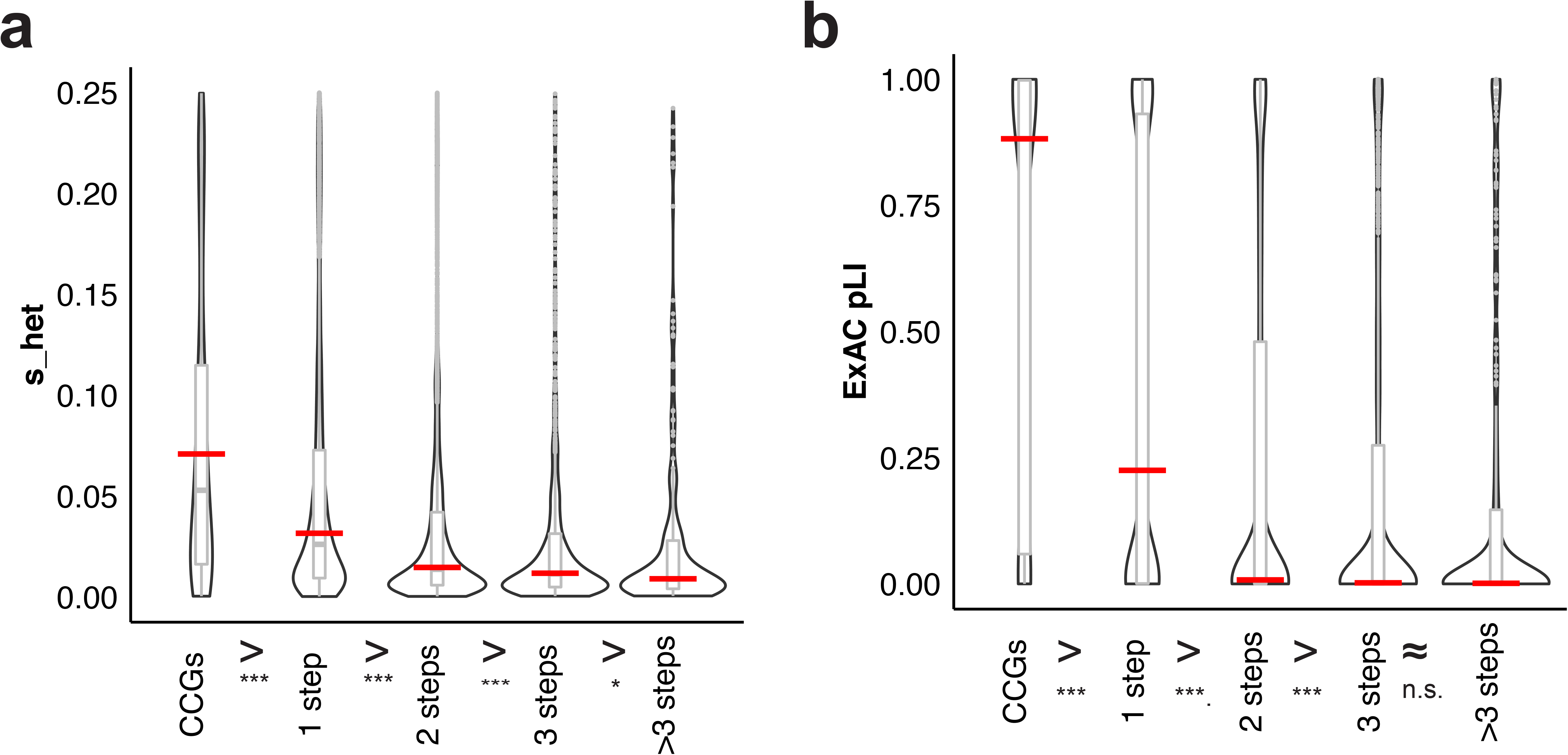
Germline selection pressure on genes in different cancer-gene neighborhood categories. **a** Selection pressure against protein-truncating variants (PTV): the lower the *S*_*h*_ score, the more tolerant the gene is for a germline PTV. **b**Loss-of-function variant intolerance (pLI): the lower the pLI score, the more tolerant the genes is for germline loss of function variants. One-sided Mann–Whitney *U* test (values of closer neighborhood genes are greater than that of all the genes in the remoter steps) *P* values are symbolized by ***, **, and * corresponding to *P* < 0.0001, *P* < 0.001, and *P* < 0.01, respectively, and n.s. abbreviating not significant. Red bars correspond to the median of the distributions. CCGs: core cancer genes.

## Discussion

Transformation from a normal cell to a cancer cell requires dysfunctions of many genes involved in key biologic process. Increasing evidence suggests that alterations in a few canonical cancer driver genes are not enough for cancer development(6, 23). To comprehensively investigate human genes that might be potentially involved in cancer development, we collected 468 CCGs from the MSK-IMPACT(13) and classified the rest of human genes into 4 categories according to their physical distance from CCGs in protein-protein interaction database. We have shown that most human genes are only 1 or 2 steps removed from canonical cancer genes in a PPI network. Genes closest to canonical cancer genes also have higher general connectivity. We demonstrated via three independent methods that cancer-biology relevant functional importance of human genes is proportional to their distance from canonical cancer genes in the network. Genes closer to cancer genes in the network have greater importance in maintaining cell viability, are under greater negative germline selection pressure, have higher somatic mutation frequency, and variants within those genes have higher cancer effect. We provide putative cancer relevance annotation for 16,932 human genes (Supplementary Table 1), this list is can help prioritizing novel genes for further functional studies in cancer research. We observed overall decreasing trend of functional importance from CCGs to neighborhood genes, but for any individual gene, its role in specific cancer type can differ substantially. Genes in each neighborhood category also have a wide range of dependency score, mutation prevalence, and effect size, which do not always correlate with one another. Some genes may have high DepMap dependency score (>0) but low mutation prevalence and effect size. Several reasons account for the variable and often discordant scores at gene level. The functional importance of most genes, even in the malignant transformation process, is likely tissue specific (24). However, there is no agreed upon list of tissue-specific cancer genes. Protein function may be affected through multiple mechanisms other than somatic mutations (transcriptional regulation, posttranslational modifications, protein degradation, binding partners, etc..). The importance of any mutation and protein dysfunction is also molecular context dependent, which in a cancer cell with unstable genome opens opportunities for a large number of potential systems level combinatorial abnormalities(24, 25). In our analysis, to maximize sample size we pooled all genes in a given neighborhood bin. This increases the statistical power for testing the general concept of neighborhood distance-dependent functional importance but could diminish the effect size itself.

## Conclusions

Overall, our findings were highly consistent across four distinct surrogates of functional importance including in vitro cancer cell viability scores after gene silencing, somatic mutation frequencies and cancer effect sizes in human cancers, and negative germline selection pressure. These results suggest that that the universe of cancer relevant genes is substantially broader than previously thought and extends to several thousand genes that can contribute functional effects when dysregulated. Neighborhood position, gene connectivity and the other functional metrics are quantitative indicators for prioritizing novel genes for further functional studies to more completely understand cancer biology.

## Methods

### Data Sources and Preparation

#### Protein-protein interactions

STRING (v11.0) is a comprehensive database of protein-protein associations using data from genomic context, high-throughput experiments, conserved co-expression and experimental results as well as text mining of the scientific literature (15). The data is available through https://string-db.org/.

#### Cancer Dependency Map (DepMap) data

DepMap project provides a gene dependency score for the majority of known human genes that represents the effect of gene silencing on cancer cell viability (17). The data is available at https://depmap.org/.

#### Somatic mutation data

Somatic mutations of 32 cancer types of 10,208 cancers in TCGA were obtained from the Multi-Center Mutation Calling in Multiple Cancers (MC3) dataset (26) that is available at https://gdc.cancer.gov/about-data/publications/mc3-2017.

#### *S*_*h*_ and pIL scores

The *S_h_* coefficients were derived from analyses of exome sequence data from 60,706 individuals and measure genome-wide estimates of selection against germline heterozygous protein-truncating variants of a gene using Bayesian estimates (20). The coefficients are available at http://genetics.bwh.harvard.edu/genescores/. The probability of being loss-of-function (LoF) intolerant (pLI) in the germline score was derived from whole exome sequence data of 60,706 individuals generated as part of the Exome-Aggregation Consortium (22) and is available at https://gnomad.broadinstitute.org/.

#### CancerMine

A text-mining based, regularly updated database of cancer driver genes, oncogenes and tumor suppressors in different types of cancer (16). Data are available at http://bionlp.bcgsc.ca/cancermine.

#### Cancer incidence rate

We obtained estimates of cancer incidence for 32 cancer types from literatures (Supplementary Table 2).

### Defining cancer gene neighbors

The shortest distance from one protein to the other in the STRING (v11.0) network was calculated by Dijkstra’s algorithm (27) using the NetworkX v1.11 Python package https://networkx.github.io/documentation/networkx-1.11/. We visualized the connection of CCGs to neighbor genes using Cytoscape (v3.7.2) with default setting (28). To plot the results (**Figure 2a**), we manually set the size of gene nodes to 50, 40, 30, 20 and 10 for CCGs, 1-step, 2-step, 3-steps and >3-step removed genes, respectively.

### Somatic mutation analysis

For somatic mutation frequency, we only considered the 2,257,845 nonsynonymous mutations that comprised missense, non-sense, frameshifting, in-frame shifting, or splice-site altering single-nucleotide changes or indels in 32 cancer types. Somatic mutation frequency at gene level was defined as the percent of cases that carried at least one nonsynonymous mutation of the gene within a cancer type. Gene level mutation frequencies were averaged over each gene neighborhood class. Tumor mutation burden (TMB) was calculated for each cancer as the number of somatic mutations, including both nonsynonymous and synonymous, per sequenced megabase. For each cancer type, we averaged TMB across all patients. The total number of genes which were affected by at least one nonsynonymous mutation was also calculated for each cancer and was averaged across all the patients with a given cancer type.

Cancer effect size is the scaled selection coefficient of the mutation, conveying the degree to which the mutation enhances the survival or reproduction of the mutant lineage. Cancer effect sizes were calculated with cancereffectsizeR 0.1.1.9006 (https://github.com/Townsend-Lab-Yale/cancereffectsizeR) as in Cannataro et al.(11) except that the likelihood of the scaled selection coefficient was maximized based on tumor-specific mutation rates, and only COSMIC v3 signatures consistent with Alexandrov et al(29) were used for each tumor type. We calculated average cancer effect size for all somatic mutations in TCGA cancer types effecting all genes in a given neighborhood category.

### Statistical analysis

The connectivity score, dependency score, somatic mutation frequency, cancer effect size, *S_h_* and pIL score were compared between different groups of genes (e.g. CCG, 1 step, 2 step, etc..) using the one-sided Mann–Whitney *U* test with the “base” package of the R-project (www.R-project.org/).

We estimated the statistical significance of the trend of the average dependency score, somatic mutation frequency, *S*_*h*_ and pIL score across the different gene groups (e.g. CCG, 1-step, 2-step, etc.) using Jonckheere Terpstra (JT) trend analysis (30). *P-*values were calculated using the “JonckheereTerpstraTest” function of “DescTools” packages (31) in the R-project. The number of permutations for the reference distribution was set as 100,000. Z statistic of Kendall’s tau (*τ*) coefficient was estimated to show the increasing (positive value) or decreasing (negative value) trend for each trend analysis. The Kendall's *τ* and z statistic were calculated using the “cor.test” function of “stats” package in the R-project (www.R-project.org/). For somatic mutation frequency, “Holm” method was used for the correction of multiple testing (32).

We separated the 32 TCGA cancer types into two groups: (i) “withTrend” indicating statistically significant decreasing trend of somatic mutation frequency, and (ii) “noTrend” corresponding to cancers with no decreasing trend. We assigned a cancer type to the withTrend group if the Jonckheere Terpstra FDR was less than 0.05, otherwise, a cancer type was assigned to the noTrend group.

We used Kendall’s *τ* to quantify association between cancer incidence rate, tumor mutation burden, and the number of affected genes. Pearson correlation coefficient and *P* value were also calculated using the “cor.test” function of “stats” package in the R-project (www.R-project.org/).

## Availability of data and materials

Data supporting the findings of this study are available in the Article, Supplementary Information, or from the authors upon request. The NetworkX v1.11 Python package for calculate protein-protein distance is available online at https://networkx.github.io/documentation/networkx-1.11/. The cancereffectsizeR 0.1.1.9006 for calculate cancer effect size is available online at https://github.com/Townsend-Lab-Yale/cancereffectsizeR.

## Authors’ contribution

L.P. and T.Q. designed the analyses. T.Q. conducted most of the analyses. H.M. performed the protein-protein interaction and gene distance analyses. V.L.C. performed cancer effect size analyses. M.M. prepared the trend analyses. J.F prepared the cancer incidence and carcinogen exposure data. L.P. and T.Q. wrote the article. M.R., J.F., H.M., M.M., J.P.T., Y.K., and M.G. contributed to data interpretation and manuscript writing.

## Acknowledgements

We thank Laurene Goode for providing clerical assistance to coordinate this project.

## References

1. Bailey MH, Tokheim C, Porta-Pardo E, Sengupta S, Bertrand D, Weerasinghe A, et al. Comprehensive Characterization of Cancer Driver Genes and Mutations. Cell. 2018;174(4):1034–5.

2. Martin GS. The hunting of the Src. Nat Rev Mol Cell Biol. 2001;2(6):467–75.

3. Sondka Z, Bamford S, Cole CG, Ward SA, Dunham I, Forbes SA. The COSMIC Cancer Gene Census: describing genetic dysfunction across all human cancers. Nat Rev Cancer. 2018;18(11):696–705.

4. Consortium ITP-CAoWG. Pan-cancer analysis of whole genomes. Nature. 2020;578(7793):82–93.

5. Hanahan D, Weinberg RA. Hallmarks of cancer: the next generation. Cell. 2011;144(5):646–74.

6. Loganathan SK, Schleicher K, Malik A, Quevedo R, Langille E, Teng K, et al. Rare driver mutations in head and neck squamous cell carcinomas converge on NOTCH signaling. Science. 2020;367(6483):1264–9.

7. Tokheim CJ, Papadopoulos N, Kinzler KW, Vogelstein B, Karchin R. Evaluating the evaluation of cancer driver genes. Proc Natl Acad Sci U S A. 2016;113(50):14330–5.

8. Iranzo J, Martincorena I, Koonin EV. Cancer-mutation network and the number and specificity of driver mutations. Proc Natl Acad Sci U S A. 2018;115(26):E6010–E9.

9. Cannataro VL, Townsend JP. Neutral Theory and the Somatic Evolution of Cancer. Mol Biol Evol. 2018;35(6):1308–15.

10. Agarwal D, Nowak C, Zhang NR, Pusztai L, Hatzis C. Functional germline variants as potential co-oncogenes. Npj Breast Cancer. 2017;3.

11. Cannataro VL, Gaffney SG, Townsend JP. Effect Sizes of Somatic Mutations in Cancer. J Natl Cancer Inst. 2018;110(11):1171–7.

12. Qing T, Mohsen H, Marczyk M, Ye YX, O’Meara T, Zhao HY, et al. Germline variant burden in cancer genes correlates with age at diagnosis and somatic mutation burden. Nat Commun. 2020;11(1).

13. Cheng DT, Mitchell TN, Zehir A, Shah RH, Benayed R, Syed A, et al. Memorial Sloan Kettering-Integrated Mutation Profiling of Actionable Cancer Targets (MSK-IMPACT): A Hybridization Capture-Based Next-Generation Sequencing Clinical Assay for Solid Tumor Molecular Oncology. J Mol Diagn. 2015;17(3):251–64.

14. Hyman DM, Solit DB, Arcilas ME, Cheng DT, Sabbatini P, Baselga J, et al. Precision medicine at Memorial Sloan Kettering Cancer Center: clinical next-generation sequencing enabling next-generation targeted therapy trials. Drug Discov Today. 2015;20(12):1422–8.

15. Szklarczyk D, Gable AL, Lyon D, Junge A, Wyder S, Huerta-Cepas J, et al. STRING v11: protein-protein association networks with increased coverage, supporting functional discovery in genome-wide experimental datasets. Nucleic Acids Res. 2019;47(D1):D607–D13.

16. Lever J, Zhao EY, Grewal J, Jones MR, Jones SJM. CancerMine: a literature-mined resource for drivers, oncogenes and tumor suppressors in cancer. Nat Methods. 2019;16(6):505–7.

17. Tsherniak A, Vazquez F, Montgomery PG, Weir BA, Kryukov G, Cowley GS, et al. Defining a Cancer Dependency Map. Cell. 2017;170(3):564–76 e16.

18. Martincorena I, Raine KM, Gerstung M, Dawson KJ, Haase K, Van Loo P, et al. Universal Patterns of Selection in Cancer and Somatic Tissues. Cell. 2017;171(5):1029–41 e21.

19. Hughes AL. Near neutrality: leading edge of the neutral theory of molecular evolution. Ann N Y Acad Sci. 2008;1133:162–79.

20. Cassa CA, Weghorn D, Balick DJ, Jordan DM, Nusinow D, Samocha KE, et al. Estimating the selective effects of heterozygous protein-truncating variants from human exome data. Nat Genet. 2017;49(5):806–10.

21. Van Hout C, Tachmazidou I, Backman J, Hoffman J, Ye B, Pandey A, et al. Whole exome sequencing and characterization of coding variation in 49,960 individuals in the UK Biobank. Eur J Hum Genet. 2019;27:1166–7.

22. Lek M, Karczewski KJ, Minikel EV, Samocha KE, Banks E, Fennell T, et al. Analysis of protein-coding genetic variation in 60,706 humans. Nature. 2016;536(7616):285–91.

23. Martincorena I, Fowler JC, Wabik A, Lawson ARJ, Abascal F, Hall MWJ, et al. Somatic mutant clones colonize the human esophagus with age. Science. 2018;362(6417):911–7.

24. Haigis KM, Cichowski K, Elledge SJ. Tissue-specificity in cancer: The rule, not the exception. Science. 2019;363(6432):1150–1.

25. Schneider G, Schmidt-Supprian M, Rad R, Saur D. Tissue-specific tumorigenesis: context matters. Nat Rev Cancer. 2017;17(4):239–53.

26. Ellrott K, Bailey MH, Saksena G, Covington KR, Kandoth C, Stewart C, et al. Scalable Open Science Approach for Mutation Calling of Tumor Exomes Using Multiple Genomic Pipelines. Cell Syst. 2018;6(3):271–81 e7.

27. Dijkstra EW. A note on two problems in connexion with graphs. Numerische Mathematik. 1959;1:269–71.

28. Shannon P, Markiel A, Ozier O, Baliga NS, Wang JT, Ramage D, et al. Cytoscape: a software environment for integrated models of biomolecular interaction networks. Genome Res. 2003;13(11):2498–504.

29. Alexandrov LB, Kim J, Haradhvala NJ, Huang MN, Tian Ng AW, Wu Y, et al. The repertoire of mutational signatures in human cancer. Nature. 2020;578(7793):94–101.

30. r JA. A distribution-free k-sample test against ordered alternatives. Biometrika. 1954(41):133–45.

31. Signorell A. DescTools: Tools for Descriptive Statistics. 2020.

32. Holm S. A Simple Sequentially Rejective Multiple Test Procedure. Scand J Stat. 1979;6(2):65–70.

