## Supplementary figures and tables for "Cancer relevance of human genes"

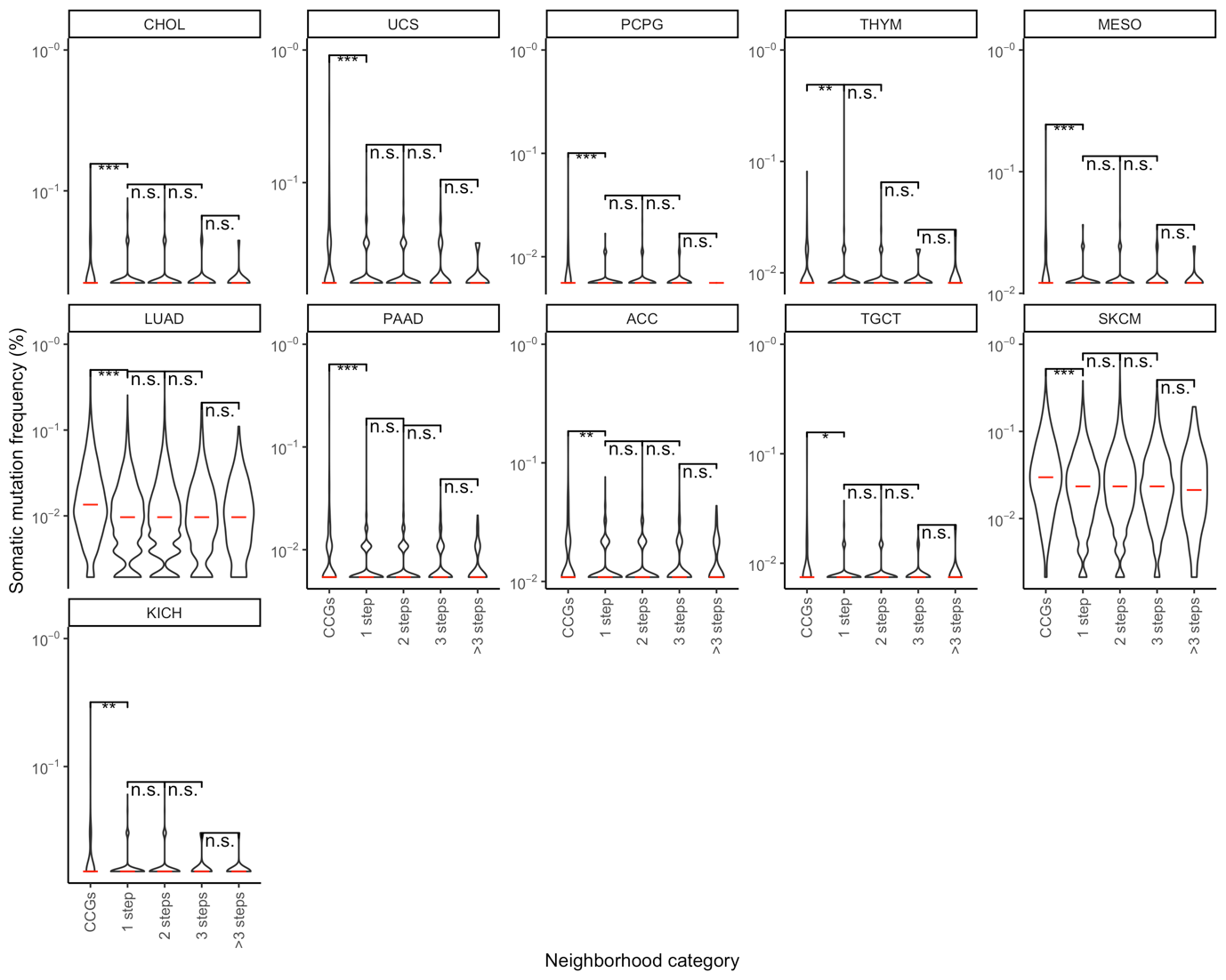

**Supplementary Figure 1. Distributions of somatic mutation frequencies of genes in different cancer gene neighborhood categories in 11 cancer types in TCGA that showed no trend by distance from CCGs.** Y-axes indicate the somatic mutation frequency. One-sided Mann–Whitney U test (values of closer neighborhood genes are greater than that of all the genes in the remoter steps) *P*-values are symbolized by ***, **, * corresponding to *P* < 0.0001, 0.001, and 0.01, respectively, n.s. = not significant. Red bars correspond to the medians of the distributions. CCGs = core cancer genes

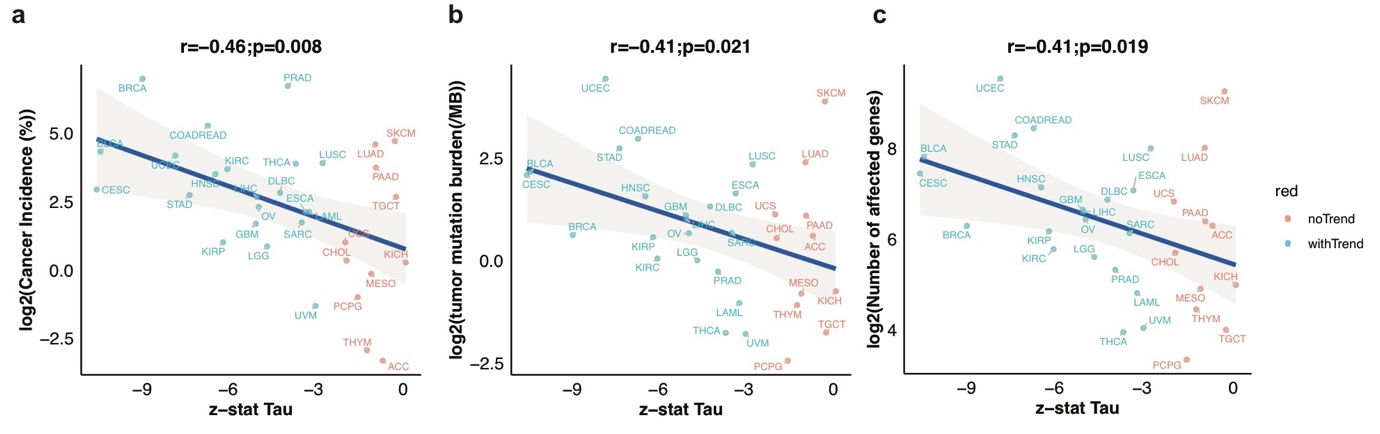

**Supplementary Figure 2. Associations between somatic mutation frequency trends and cancer incidence, tumor mutation burden (TMB), and number of mutated genes across all TCGA cancer types.** (**a**) Z-statistic τ versus cancer incidence rate. (**b**) Z-statistic τ versus TMB. (**c**) Z-statistic τ versus number of affected genes by somatic mutation. Each dot represents a cancer type and the colors indicate if it showed a significant trend for decreasing average somatic mutation frequency across CCG neighborhood categories with increasing distance from core cancer genes. The Y-axes shows average values within cancer types. The r represents Pearson correlation coefficient. *P* values was estimated based on the Pearson product-moment correlation coefficient.

**Supplementary Table 1. Trend analysis of somatic mutation frequencies in TCGA cancer types**

|  | **tissues** | **JT Statistic** | **JT Pvalue** | **JT FDR** | **Kendall's Tua**  **z-stat** | **Group** |
| --- | --- | --- | --- | --- | --- | --- |
| 1 | CESC | 22807252.5 | 1.00E-05 | 0.00033 | -10.87633073 | withTrend |
| 2 | BLCA | 27667467.5 | 1.00E-05 | 0.00033 | -10.74287481 | withTrend |
| 3 | BRCA | 26938261 | 1.00E-05 | 0.00033 | -9.268878675 | withTrend |
| 4 | UCEC | 34463028.5 | 1.00E-05 | 0.00033 | -8.12041567 | withTrend |
| 5 | STAD | 30550646.5 | 1.00E-05 | 0.00033 | -7.626321569 | withTrend |
| 6 | COADREAD | 33294429 | 1.00E-05 | 0.00033 | -6.982576229 | withTrend |
| 7 | HNSC | 25655920 | 1.00E-05 | 0.00033 | -6.721988932 | withTrend |
| 8 | KIRP | 11071585.5 | 1.00E-05 | 0.00033 | -6.450495652 | withTrend |
| 9 | KIRC | 10824987 | 1.00E-05 | 0.00033 | -6.300903146 | withTrend |
| 10 | GBM | 20445726.5 | 1.00E-05 | 0.00033 | -5.304454872 | withTrend |
| 11 | LIHC | 17712084.5 | 1.00E-05 | 0.00033 | -5.273644225 | withTrend |
| 12 | OV | 16873215.5 | 1.00E-05 | 0.00033 | -5.20168939 | withTrend |
| 13 | LGG | 14580339 | 1.00E-05 | 0.00033 | -4.90506384 | withTrend |
| 14 | DLBC | 1102995 | 1.00E-05 | 0.00033 | -4.45346198 | withTrend |
| 15 | PRAD | 11357893 | 2.00E-05 | 0.00036 | -4.183073424 | withTrend |
| 16 | THCA | 2917159 | 5.00E-05 | 0.00085 | -3.907338911 | withTrend |
| 17 | SARC | 8348295.5 | 0.00015 | 0.0024 | -3.695840208 | withTrend |
| 18 | ESCA | 12512715.5 | 0.00023 | 0.00345 | -3.564258986 | withTrend |
| 19 | LAML | 886884 | 0.00035 | 0.0049 | -3.435109273 | withTrend |
| 20 | UVM | 139256 | 6.00E-04 | 0.0078 | -3.214400171 | withTrend |
| 21 | LUSC | 30123906.5 | 0.00151 | 0.01812 | -2.967343395 | withTrend |
| 22 | UCS | 2865321 | 0.01523 | 0.16753 | -2.172671635 | noTrend |
| 23 | CHOL | 305588.5 | 0.01674 | 0.16753 | -2.126036614 | noTrend |
| 24 | PCPG | 288630.5 | 0.0395 | 0.3555 | -1.736827249 | noTrend |
| 25 | THYM | 590480 | 0.07889 | 0.63112 | -1.408916469 | noTrend |
| 26 | MESO | 512302.5 | 0.1028 | 0.7196 | -1.260442688 | noTrend |
| 27 | LUAD | 30446669.5 | 0.13273 | 0.79638 | -1.118644598 | noTrend |
| 28 | PAAD | 11000464 | 0.13746 | 0.79638 | -1.096141153 | noTrend |
| 29 | ACC | 3045342 | 0.19608 | 0.79638 | -0.857514619 | noTrend |
| 30 | SKCM | 34638426.5 | 0.33058 | 0.99174 | -0.437186104 | noTrend |
| 31 | TGCT | 362331 | 0.34681 | 0.99174 | -0.394557849 | noTrend |
| 32 | KICH | 393465 | 0.47669 | 0.99174 | -0.054696354 | noTrend |

|  | **Cancer type** | **Abbreviation** | **Incidence Rate**  **(per 100,000)** |
| --- | --- | --- | --- |
| 1 | Breast invasive carcinoma | BRCA | 125 |
| 2 | Prostate adenocarcinoma | PRAD | 104.1 |
| 3 | Skin cutaneous melanoma | SKCM | 25.8 |
| 4 | Lung adenocarcinoma | LUAD | 23.7 |
| 5 | Bladder urothelial carcinoma | BLCA | 19.8 |
| 6 | Colon Rectal adenocarcinoma | COADREAD | 38.2 |
| 7 | Uterine corpus endometrial carcinoma | UCEC | 17.9 |
| 8 | Lung squamous cell carcinoma | LUSC | 14.8 |
| 9 | Thyroid carcinoma | THCA | 14.6 |
| 10 | Pancreatic adenocarcinoma | PAAD | 13.2 |
| 11 | Renal clear cell carcinoma | KIRC | 12.7 |
| 12 | Head and neck squamous cell carcinoma | HNSC | 11.2 |
| 13 | Cervical squamous cell carcinoma and endocervical adenocarcinoma | CESC | 7.6 |
| 14 | Lymphoid Neoplasm Diffuse Large B-cell Lymphoma | DLBC | 7 |
| 15 | Stomach adenoarcinoma | STAD | 6.6 |
| 16 | Testicular germ cell tumors | TGCT | 6.3 |
| 17 | Hepatocellular carcinoma | LIHC | 6.3 |
| 18 | Ovarian serous cystadenocarcinoma | OV | 4.9 |
| 19 | Esophageal carcinoma | ESCA | 4.3 |
| 20 | Acute Myeloid Leukemia | LAML | 4.3 |
| 21 | Sarcoma (soft tissue) | SARC | 3.3 |
| 22 | Glioblastoma | GBM | 3.2 |
| 23 | Renal papillary cell carcinoma | KIRP | 2 |
| 24 | Uterine carcinosarcoma | UCS | 2 |
| 25 | Brain lower grade glioma | LGG | 1.8 |
| 26 | Cholangiocarcinoma | CHOL | 1.26 |
| 27 | Renal chromophobe | KICH | 1.2 |
| 28 | Mesothelioma | MESO | 0.9 |
| 29 | Pheochromocytoma and Paraganglioma | PCPG | 0.5 |
| 30 | Uveal melanoma | UVM | 0.4 |
| 31 | Thymoma | THYM | 0.13 |
| 32 | Adrenocortical carcinoma | ACC | 0.1 |

**Supplementary Table 2. Cancer incidence rate.**
